# A PDZ-kinase allosteric relay mediates Par complex regulator exchange

**DOI:** 10.1101/2024.10.18.619144

**Authors:** Elizabeth Vargas, Rhiannon R. Penkert, Kenneth E. Prehoda

## Abstract

The Par complex polarizes the plasma membrane of diverse animal cells using the catalytic activity of atypical Protein Kinase C (aPKC) to pattern substrates. Two upstream regulators of the Par complex, Cdc42 and Par-3, bind separately to the complex to influence its activity in different ways. Each regulator binds a distinct member of the complex, Cdc42 to Par-6 and Par-3 to aPKC, making it unclear how they influence one another’s binding. Here we report the discovery that Par-3 binding to aPKC is regulated by aPKC autoinhibition and link this regulation to Cdc42 and Par-3 exchange. The Par-6 PDZ domain activates aPKC binding to Par-3 via a novel interaction with the aPKC kinase domain. Cdc42 and Par-3 have opposite effects on the Par-6 PDZ–aPKC kinase interaction: while the Par-6 kinase domain interaction competes with Cdc42 binding to the complex, Par-3 binding is enhanced by the interaction. The differential effect of Par-3 and Cdc42 on the Par-6 PDZ interaction with the aPKC kinase domain forms an allosteric relay that connects their binding sites and is responsible for the negative cooperativity that underlies Par complex polarization and activity.

## Introduction

The Par complex plays a central role in establishing and maintaining cortical polarity in a wide variety of animal cells, including epithelial, immune and neural stem cells (1–5). The complex is composed of the proteins atypical kinase C (aPKC) and Par-6, with the catalytic activity of aPKC providing the primary output of the complex (6–8). Several upstream regulators bind directly to the Par complex to precisely control its localization and activity. Regulators include the Rho GTPase Cdc42, which is thought to activate aPKC activity, and the multi-PDZ protein Par-3 (Bazooka or Baz in `Drosophila`), which is thought to repress activity (9–12). The opposing effects of the Par regulators are facilitated by negative cooperativity that ensures only one of the regulators is bound to the complex at a time (13). Given that each regulator binds to a different member of the Par complex, Cdc42 to Par-6 (14–18) and Par-3 to aPKC (19–22), it has been unclear how they might influence one another’s binding. Here we examine the mechanism by which negative cooperativity is implemented in the Par complex to support the distinct activities induced by Cdc42 and Par-3.

The combined regulation by Cdc42 and Par-3 ensure that the Par complex is activated at the proper time and in the appropriate membrane domain, and the presence of both regulators is generally required for Par-mediated polarity in a wide array of systems (23–28). In the `Drosophila` neuroblast, both Cdc42 and Baz are apically enriched during asymmetric cell division and disruption of either disrupts localization of the Par complex (10, 29). In `cdc42` mutants, Par-6 and aPKC are both found in the cytoplasm, even though Baz is still apically enriched, suggesting the regulators work independently and sequentially to properly localize the Par complex (10). In the `C. elegans` zygote, Par-3 maintains the Par complex in an inactive state while coupling it to actomyosin-generated cortical flows which moves the complex toward the anterior cortex (30, 31). At the anterior cortex, GTP-bound Cdc42 stimulates aPKC activity. The localization of the Par complex in `C. elegans` is consistent with distinct Par-3-bound and Cdc42-bound pools, suggesting that the Par complex switches between regulator bound states (9). This exchange between Cdc42 and Par-3-bound states can be recapitulated `in vitro` with purified components demonstrating that negative cooperativity underlies complex switching and that no additional factors are required for switching (13).

Cdc42 and Par-3 bind distinct sites on the Par complex making it likely that an allosteric mechanism underlies their competitive binding. Par-6 contains a CRIB motif that selectively binds GTP-bound Cdc42 (Fig. 1A) (14, 15, 18). Upon binding, Cdc42 induces an allosteric transition in the adjacent Par-6 PDZ domain which influences the PDZ’s affinity for the transmembrane receptor Crumbs (32). The kinase domain and an adjacent PDZ Binding Motif (PBM) of aPKC primarily interact with the second of three Par-3 PDZ domains (19), though other interactions have been reported (14, 15, 33–35). The only reported biochemical interaction between aPKC and Par-6 is through their PB1 domains (36, 37), though the Par-6 PDZ domain has been proposed to inhibit the aPKC kinase domain activity via an unknown mechanism (38). Using purified components in an affinity pulldown assay, we were able to identify the elements of the Par complex that are required for Cdc42 and Par-3 complex switching. Here we uncover a direct interaction between the PDZ domain of Par-6 and the kinase domain of aPKC that allows Cdc42 to toggle the affinity of the Par complex for Par-3, providing a mechanism for how complex switching may occur `in vivo`.

**Figure 1:**
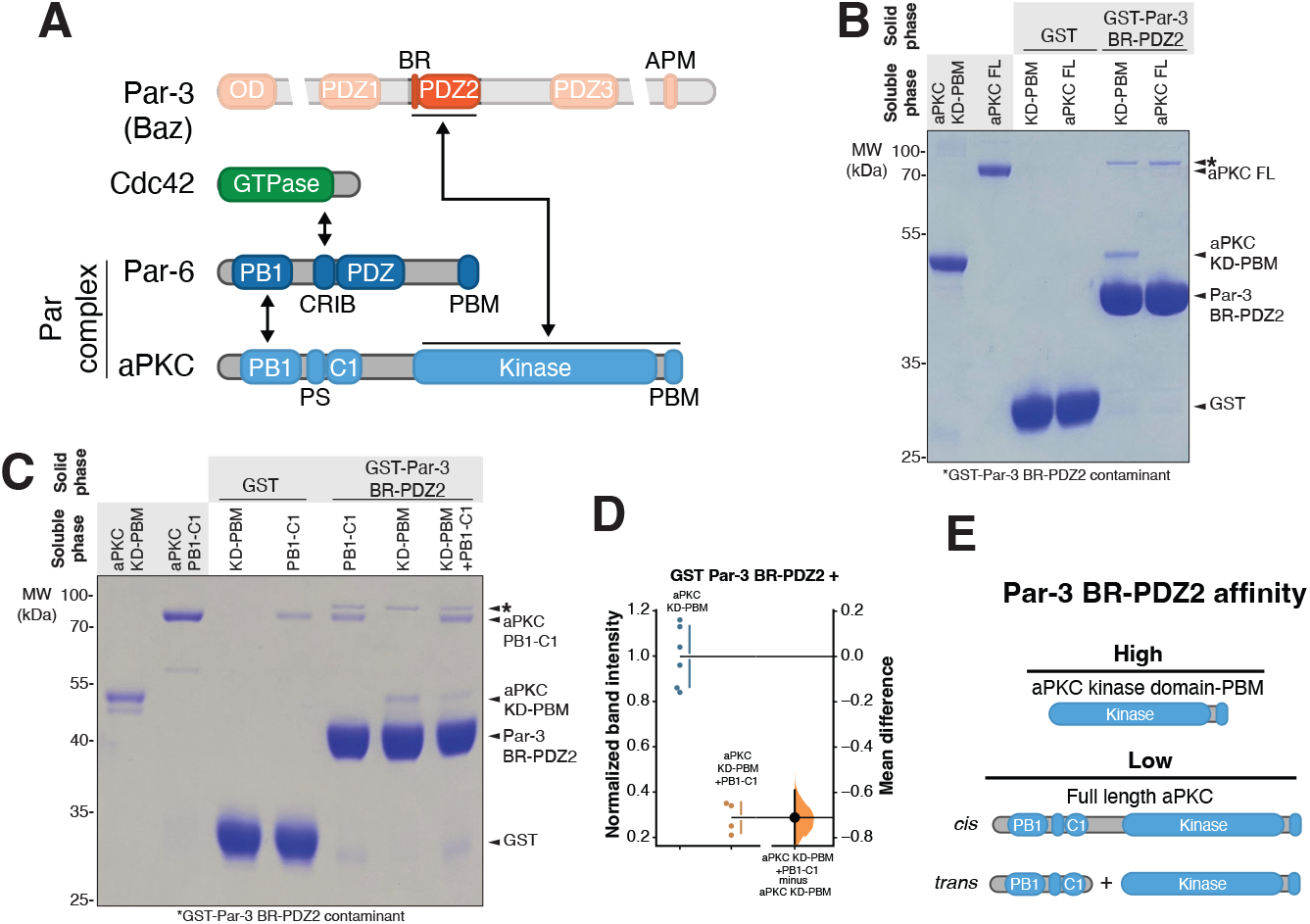
Par-3 binding to aPKC is autoinhibited by the regulatory domain of aPKC. **A**.Schematic of the domain architecture of and interactions between the Par complex proteins (aPKC and Par-6) and known regulators Par-3 (Bazooka in *Drosophila*) and Cdc42. **B**.Par-3 binding to aPKC kinase domain with its associated PDZ-binding motif (KD-PBM) or full length aPKC (aPKC FL). Solid phase (glutathione resin)-bound glutathione-S-transferase (GST) or GST-fused Par-3 PDZ2 with its associated basic region (BR-PDZ2) incubated with aPKC KD-PBM or full length (aPKC FL). *Shaded regions* indicate the fraction applied to the gel (soluble phase or solid phase components after incubation with indicated soluble components and washing). **C**.Par-3 binding to aPKC KD-PBM and/or regulatory domain (aPKC PB1-C1). Labeling as described in (B). **D**.Gardner-Altman estimation plot of normalized band intensity of aPKC KD-PBM binding to Par-3 BR-PDZ2 in the presence or absence of its regulatory domain (aPKC PB1-C1). The results of each replicate (filled circles) are shown along with mean and standard deviation (gap and bars adjacent to filled circles). The mean difference is plotted on the right as a bootstrap sampling distribution (shaded region) with a 95% confidence interval (black error bar). **E**.Summary of Par-3 interactions with aPKC.

## Results

### Par-3 binding to aPKC is autoinhibited and activated by Par-6

The region of Par-3 containing a short basic region followed by its PDZ2 domain (BR-PDZ2; hereafter PDZ2) binds to the aPKC kinase domain and its PDZ Binding Motif (KD-PBM; Fig. 1A) and this interaction is essential for aPKC membrane recruitment and polarization (19, 20). The interaction is high affinity, with an overall ΔG° of 7.9 kcal/mol (*K*_d_ = 1.3 µM) (19). We examined Par-3 PDZ2 binding to full-length aPKC to determine if the known intramolecular interactions within aPKC influence Par-3 binding. The interaction between Par-3 PDZ2 and full length aPKC was significantly reduced compared to the KD-PBM alone (Fig. 1B), suggesting that Par-3’s interaction with the KD-PBM is repressed when the regulatory module (comprised of the PB1, PS and C1 domains) is included (i.e. aPKC is autoinhibited with respect to Par-3 binding). We also observed inhibition of Par-3 PDZ2 binding to the KD-PBM when the aPKC regulatory module is added `in trans` (Fig. 1C-D), confirming that it interferes with Par-3 binding to aPKC. Thus, in addition to its established role in regulating catalytic activity (39), autoinhibition of aPKC regulates binding to Par-3.

Autoinhibition of Par-3 binding to aPKC raises the question of how binding to Par-3 becomes activated. Previously, we found that the Par complex, which consists of aPKC bound to Par-6, binds to Par-3 PDZ2 with a similar affinity (ΔG° of 7.5 kcal/mol; Kd = 2.5 µM) to aPKC KD-PBM alone suggesting that full-length Par-6 activates aPKC’s ability to bind Par-3 (19). When we compared the binding of Par complex to Par-3 with the binding of full length aPKC to Par-3, we found that the Par complex bound substantially better (∼8-fold) than full length aPKC alone to Par-3 PDZ2 (Fig. 2A-B) confirming that Par-6 overcomes aPKC autoinhibition of Par-3 binding. These results indicate that the Par complex contains a Par-3 binding site (the aPKC KD-PBM), an element that represses the Par-3 binding site (the aPKC regulatory module), and other elements within Par-6 that disrupt this repression.

**Figure 2:**
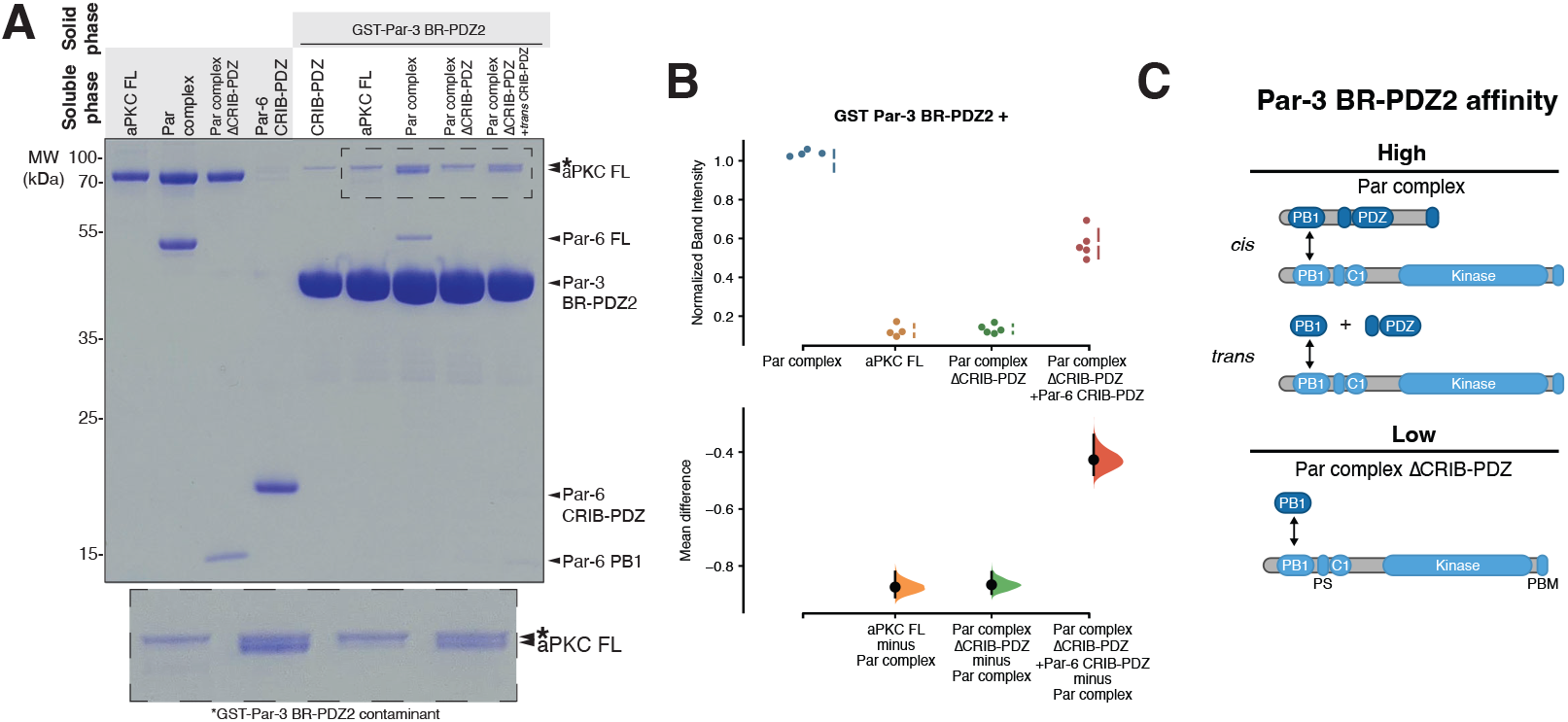
Par-3 binding to aPKC is activated by Par-6. **A**.Par-3 binding to aPKC in the presence or absence of various Par-6 domains. Solid phase (glutathione resin)-bound glutathione-S-transferase (GST)-fused Par-3 PDZ2 with its associated basic region (BR-PDZ2) incubated with full length aPKC (aPKC FL), Par complex (full length aPKC and Par-6), Par complex ΔCRIB-PDZ (aPKC with the Par-6 PB1 domain) or Par complex ΔCRIB-PDZ plus Par-6 CRIB-PDZ. *Shaded regions* indicate the fraction applied to the gel (soluble phase or solid phase components after incubation with indicated soluble components and washing). Inset shows enlargement of the last 4 lanes as indicated. **B**.Cumming estimation plot of the normalized band intensity of aPKC binding to Par-3 BR-PDZ2 under the indicated conditions shown in (A). The result of each replicate (filled circles) along with the mean and SD (gap and bars next to circles) are plotted in the top panel and the mean differences are plotted in the bottom panel as a bootstrap sampling distribution (shaded region) with a 95% confidence interval (black error bar). **C**.Summary of Par-3 interactions with Par complex.

### The Par-6 CRIB-PDZ promotes aPKC binding to Par-3

We sought to identify the Par-6 elements that activate Par-3 binding to aPKC. Par-6 interacts with aPKC via a PB1-PB1 interaction [Fig. 1A; (36, 37)] suggesting that the Par-6 PB1 could be responsible for activating aPKC’s Par-3 binding. To determine if the Par-6 PB1 domain is sufficient to activate Par-3 binding we examined the binding to full length aPKC in the presence of Par-6 PB1 (“Par complex ΔCRIB-PDZ”) and found that Par-3 PDZ2 bound with an affinity similar to that of full length aPKC alone (Fig. 2A-B). Thus, binding of the Par-6 PB1 domain is insufficient to relieve aPKC autoinhibition and facilitate high affinity Par-3 binding.

Aside from a PB1 domain, Par-6 contains a CRIB domain, which is known to bind Cdc42 (14, 15, 18), and a PDZ domain that interacts with other polarity proteins such as Stardust and Crumbs (32, 40). When Par-6 CRIB-PDZ was added to Par complex ΔCRIB-PDZ we observed enhanced binding to Par-3 (Fig 2A-B). Binding was not fully restored, likely because of effective concentration effects when the PB1 and CRIB-PDZ domains are covalently attached. Importantly, however, activation of Par-3 binding by Par-6 CRIB-PDZ `in trans` indicates that the covalent linkage between PB1 and CRIB-PDZ is not required to relieve aPKC autoinhibition.

### The Par-6 PDZ domain binds to the aPKC kinase domain

How does the Par-6 CRIB-PDZ activate aPKC binding to Par-3? No interactions have been reported between aPKC and Par-6 outside of the PB1-PB1 interaction, but our data suggest that Par-6 CRIB-PDZ may interact directly with aPKC. We tested if CRIB-PDZ could bind to either the N-terminus of aPKC, which contains the regulatory module (PB1-C1), or the C-terminus of aPKC, which contains the kinase and PBM domains (KD-PBM). We found that Par-6 CRIB-PDZ bound directly to aPKC KD-PBM, but not PB1-C1 (Fig. 3A). When we examined the interaction at higher resolution, we found that Par-6 PDZ (lacking the CRIB motif) bound aPKC KD-PBM with a similar affinity to that of Par-6 CRIB-PDZ (Fig. 3B). Interestingly, despite the interaction involving a PDZ domain, the PBM (PDZ-binding motif) of aPKC was not required, though binding may be reduced in its absence (Fig. 3B). Thus, we have discovered a new interaction between aPKC and Par-6 outside of the PB1-PB1 heterodimerization, between the aPKC KD and Par-6 PDZ.

**Figure 3:**
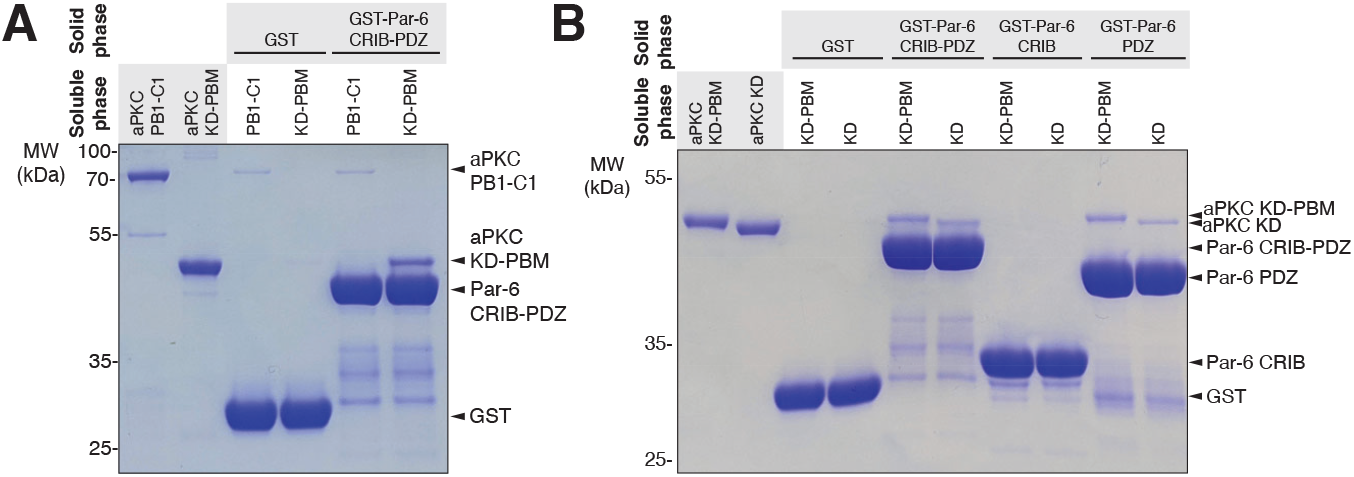
Par-6 PDZ binds directly to the kinase domain of aPKC. **A**.Par-6 CRIB-PDZ binding to the various domains of aPKC. Solid phase (glutathione resin)-bound glutathi-one-S-transferase (GST)-fused Par-6 CRIB-PDZ incubated with the regulatory module of aPKC (PB1-C1) or aPKC kinase domain (KD) with its PDZ-binding motif (PBM). *Shaded regions* indicate the fraction applied to the gel (soluble phase or solid phase components after incubation with indicated soluble components and washing). **B**.Binding of individual domains of Par-6 to aPKC KD or KD-PBM. Solid phase (glutathione resin)-bound glutathi-one-S-transferase (GST)-fused Par-6 CRIB-PDZ, GST-fused Par-6 CRIB or GST-fused Par-6 PDZ incubated with either aPKC KD-PBM or aPKC KD only. *Shaded regions* indicate the fraction applied to the gel (soluble phase or solid phase components after incubation with indicated soluble components and washing).

### Par-3 PDZ2 and Par-6 CRIB-PDZ bind cooperatively to aPKC

Given that both Par-3 PDZ2 and Par-6 PDZ appear to bind the aPKC KD, we sought to determine if they influenced one another’s binding. The presence of Par-6 CRIB-PDZ significantly enhanced Par-3 binding to full length aPKC (Figure 2) indicating positive cooperativity of the interaction. To confirm binding cooperativity, we examined if Par-3 could also enhance binding of Par-6 CRIB-PDZ to aPKC. In the absence of Par-3, Par complex ΔCRIB-PDZ only binds weakly to Par-6 CRIB-PDZ (Fig. 4A). In the presence of Par-3 PDZ2, the binding of aPKC to Par-6 CRIB-PDZ was increased greater than 20-fold (Fig. 4B) demonstrating that Par-3 and Par-6 CRIB-PDZ significantly enhance one another’s binding to aPKC KD-PBM.

**Figure 4:**
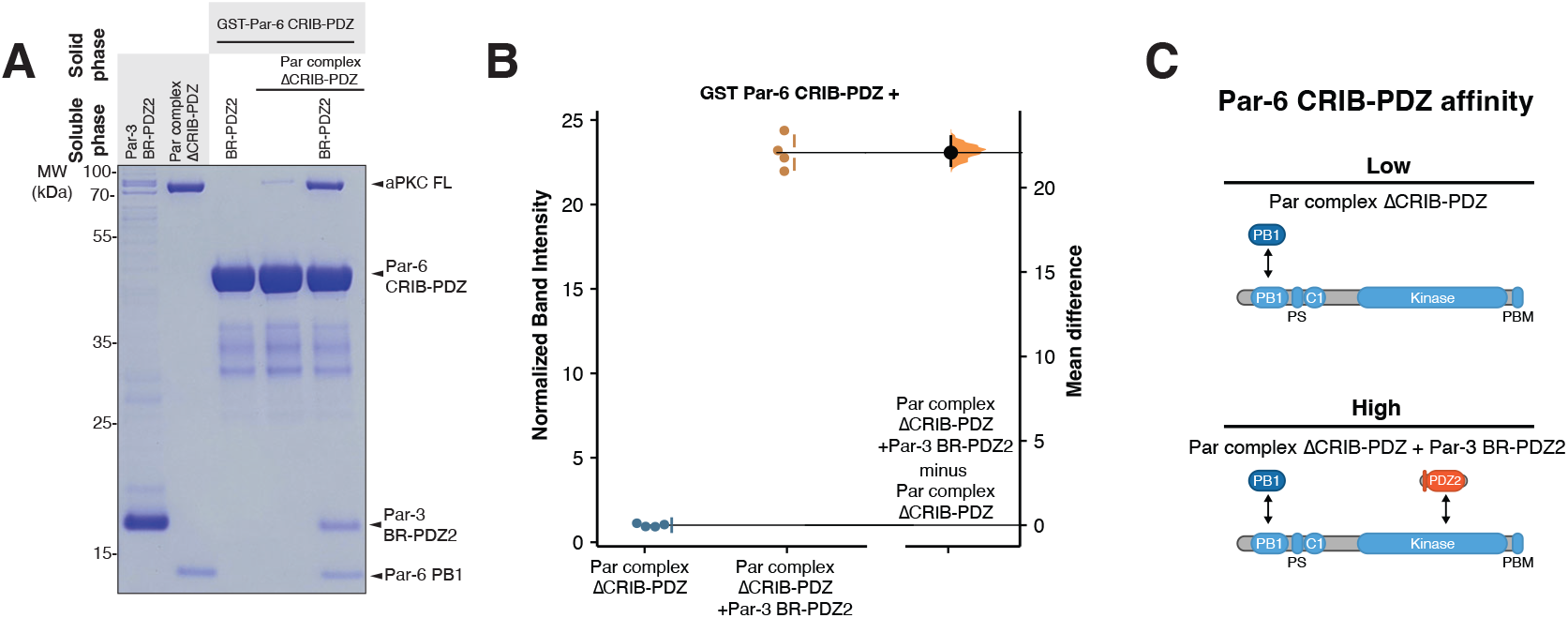
Par-6 CRIB-PDZ and Par-3 PDZ2 bind cooperatively to aPKC. **A**.Binding of Par-6 CRIB-PDZ to aPKC in the presence or absence of Par-3. Solid phase (glutathione resin)-bound glutathione-S-transferase (GST)-fused Par-6 CRIB-PDZ incubated with Par complex ΔCRIB-PDZ in the presence or absence of Par-3 PDZ2 and its associated basic region (BR-PDZ2). Shaded regions indicate the fraction applied to the gel (soluble phase or solid phase components after incubation with indicated soluble components and washing). **B**.Gardner-Altman estimation plot of normalized band intensity of aPKC binding to Par-6 CRIB-PDZ in the absence or presence of Par-3 BR-PDZ2. **C**.Summary of Par-6 CRIB-PDZ and Par-3 BR-PDZ2 cooperative binding to aPKC

### Cdc42 displaces Par-6 CRIB-PDZ from aPKC KD-PBM to regulate Par-3 binding to aPKC

The Par-6 PDZ interaction with the aPKC KD (Figure 3) and its cooperative binding with Par-3 PDZ2 (Figure 4) suggest a possible mechanism for Cdc42 and Par-3 complex switching. If Cdc42 reduces the affinity of Par-6 PDZ for aPKC KD, the positive cooperativity of Par-6 PDZ and Par-3 binding would be lost, returning aPKC to the low Par-3 affinity state. Cdc42 is known to induce an allosteric change in the Par-6 PDZ domain when it binds the CRIB motif (41), providing a possible mechanism for altering the Par-6 PDZ affinity for the aPKC KD. We directly tested the effect of constitutively active Cdc42 (Cdc42^Q61L^; hereafter Cdc42) on the interaction between Par-6 CRIB-PDZ and aPKC KD-PBM and found that Cdc42 reduced the affinity of the aPKC KD-PBM interaction with Par-6 CRIB-PDZ to an extent that it was virtually undetectable in our assay (Fig. 5A-B). Consistent with this observation, we previously found that Cdc42 has a higher affinity for Par-6 CRIB-PDZ alone (ΔG° of 7.6 kcal/mole; *K*_d_ = 2.2 µM) than for the Par complex, (ΔG° of 7.1 kcal/mole; *K*_d_ = 5.4 µM) (13). Thus, while the Par-6 CRIB-PDZ interaction with aPKC KD-PBM enhances the affinity for Par-3 (i.e. positive cooperativity), it lowers the affinity for Cdc42 (i.e. negative cooperativity). We confirmed this behavior using reconstituted Par complex with the `trans` CRIB-PDZ (Fig. 6A) by determining if Cdc42 displaced Par-3 binding in this context. We observed a significant reduction in Par-3 binding upon addition of Cdc42 along with displacement of the CRIB-PDZ (Fig. 6A-B).

**Figure 5:**
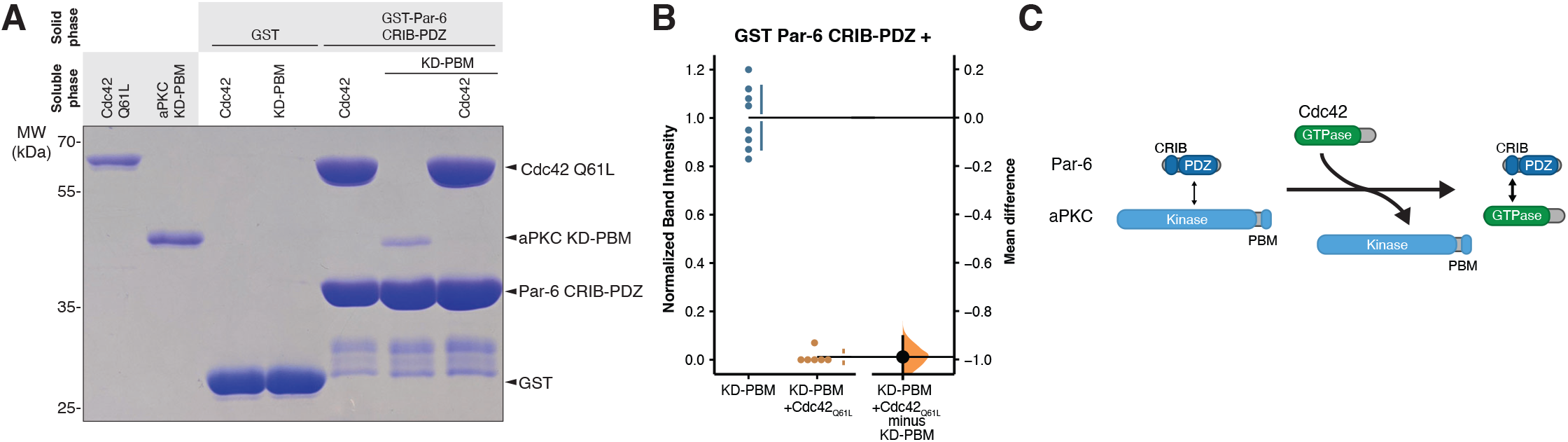
Cdc42 displaces Par-6 CRIB-PDZ from the aPKC kinase domain (KD). **A**.Binding of Par-6 CRIB-PDZ to Cdc42 and aPKC KD with its PDZ binding motif (KD-PBM). Solid phase (glutathione resin)-bound glutathione-S-transferase (GST)-fused Par-6 CRIB-PDZ incubated with Cdc42 Q61L, aPKC KD-PBM or both Cdc42 and aPKC KD-PBM. **B**.Gardner-Altman estimation plot of normalized band intensity of aPKC KD-PBM binding to Par-6 CRIB-PDZ in the absence or presence of Cdc42 Q61L. **C**.Summary of Cdc42’s effect on the Par-6 CRIB-PDZ interaction with the aPKC KD.

**Figure 6:**
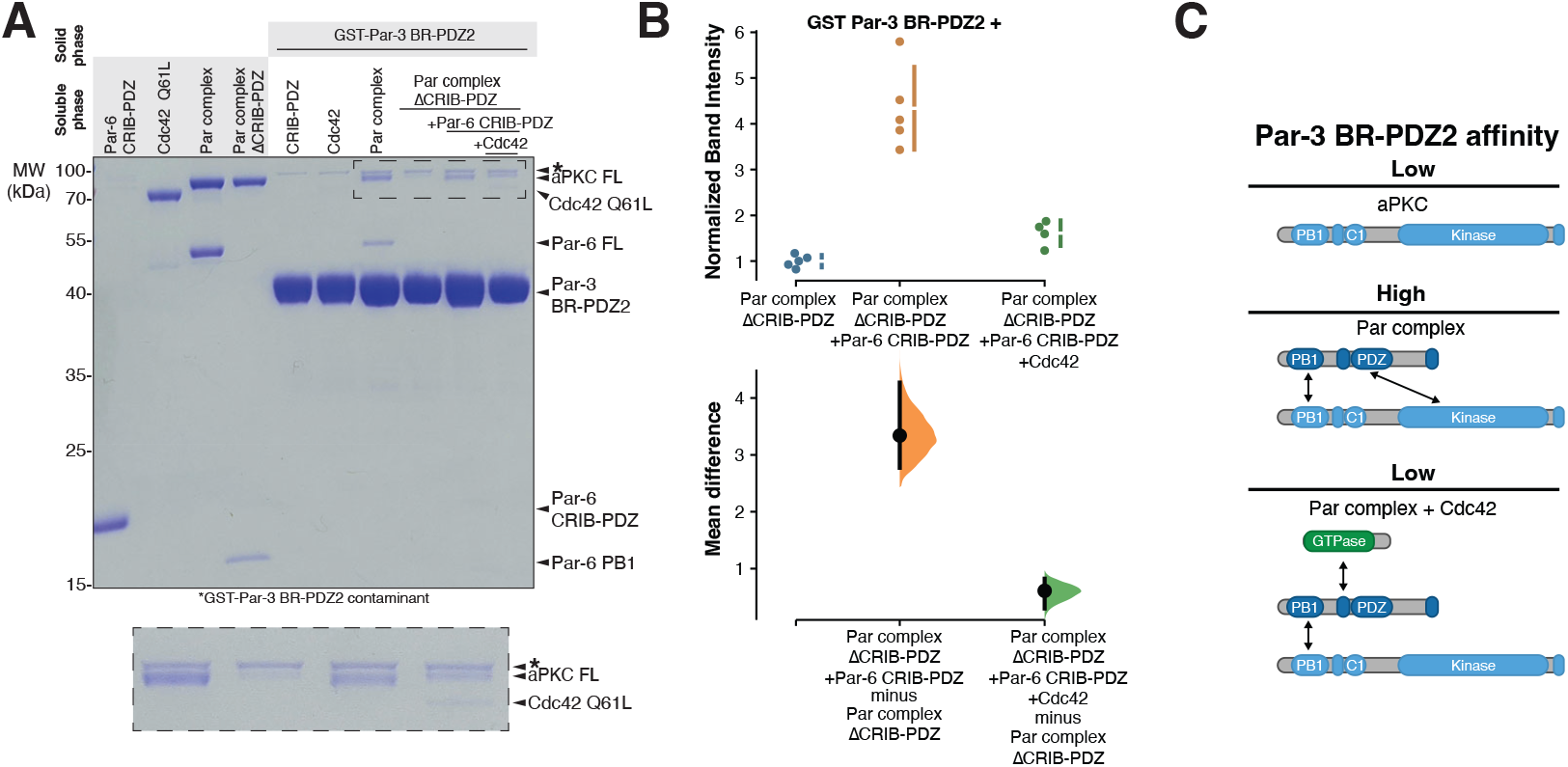
Cdc42 toggles the interaction between Par-6 CRIB-PDZ and aPKC kinase domain (KD) to influence Par-3 binding. **A**.Binding of Par-3 to aPKC in the presence of Par-6 CRIB-PDZ with or without Cdc42. Solid phase (glutathione resin)-bound glutathione-S-transferase (GST)-fused Par-3 PDZ2 with its associated basic region (BR-PDZ2) incubated with Par complex (full length aPKC and Par-6) or Par complex ΔCRIB-PDZ (full length aPKC plus the Par-6 PB1 domain) with Par-6 CRIB-PDZ and Cdc42Q61L as indicated. Shaded regions indicate the fraction applied to the gel (soluble phase or solid phase components after incubation with indicated soluble components and washing). Inset shows enlargement of the last 4 lanes. **B**.Cumming estimation plot of normalized band intensity of aPKC binding to Par-3 BR-PDZ2 under the indicated conditions from (A). The result of each replicate (filled circles) along with the mean and SD (gap and bars next to circles) are plotted in the top panel and the mean differences are plotted in the bottom panel as a bootstrap sampling distribution (shaded region) with a 95% confidence interval (black error bar). **C**.Summary of PDZ-kinase mediated regulator exchange. Par-3 BR-PDZ2 binds with low affinity to aPKC because of autoinhibition. Par-6 relieves autoinhibition through its PDZ domain interacting with the aPKC kinase domain leading to high affinity Par-3 binding. Cdc42 inhibits the PDZ-kinase interaction to restore aPKC to its low Par-3 affinity state.

## Discussion

A critical step in current models for Par-mediated polarity is the exchange between Par-3- and Cdc42-bound Par complex. These regulators promote distinct activities with their combined action resulting in a polarized, active Par complex. Par-3 maintains the complex in an inactive state while coupling it to actomyosin-driven cortical movements that polarize the complex (30, 31). Conversely, Cdc42 activates aPKC’s catalytic activity (12, 16), which is essential for polarizing substrates. Due to their differing effects on aPKC catalytic activity, Par-3 and Cdc42 form mutually exclusive interactions with the complex (9). This regulator exchange is driven by negative cooperativity in their coupled interactions with the complex (13). However, the physical features of the complex that couple Par-3 and Cdc42 binding were previously unknown. Our research has uncovered a crucial internal Par complex interaction between the Par-6 PDZ domain and the aPKC catalytic domain (KD) that plays a key role in the exchange process. Both Cdc42 binding to Par-6 CRIB-PDZ and Par-3 binding to aPKC KD-PBM are coupled to the interaction with the kinase domain. Notably, Par-3 and Cdc42 exert opposite effects on this interaction: Par-3 binding enhances it (positive cooperativity), whereas Cdc42 reduces it (negative cooperativity). These opposing actions of Par-3 and Cdc42 support a mechanism for Par-3 and Cdc42 complex exchange. The CRIB-PDZ binding to KD-PBM promotes Par-3 binding to aPKC but is detrimental to Cdc42 binding, thus facilitating the exchange between Par-3- and Cdc42-bound states of the complex.

How might Cdc42 influence the Par-6 CRIB-PDZ interaction with the aPKC KD-PBM? The CRIB and PDZ domains of Par-6 are structurally linked, with the CRIB forming an extension of a beta-sheet that runs through the PDZ domain (18). The CRIB-PDZ linkage mediates an allosteric change in the PDZ domain upon Cdc42 binding to the CRIB, changing the PDZ’s structure and its affinity for standard PDZ binding motifs (41). Our results suggest that it also alters the PDZ’s affinity for the aPKC kinase domain. Cdc42 binding to the CRIB increases the PDZ’s affinity for standard COOH-terminal PBM ligands (41), or does not influence binding of non-COOH terminal “internal” ligands (40). Here, we found that Cdc42 decreases the affinity of the PDZ for the kinase domain. The opposing effects of Cdc42 on Par-6 PDZ ligands suggest that the kinase domain binds the Par-6 PDZ elsewhere from its PBM binding site.

The biochemical properties of Par-3 PDZ2 and Par-6 PDZ suggest a mechanism for how they simultaneously interact with aPKC KD-PBM. Both the aPKC KD and PBM are required for high affinity Par-3 PDZ2 binding (19), suggesting that this PDZ forms a canonical PDZ-PBM interaction alongside contacts with the kinase domain. Par-6 PDZ binding depends primarily on the aPKC KD rather than the PBM, suggesting that it forms a non-canonical interaction, potentially on a surface outside of its PBM binding site. The Par-6 PDZ and Par-3 PDZ2 bind cooperatively to aPKC KD-PBM, suggesting that they contact one another once they form a ternary complex. Our results suggest how these interactions mediate complex exchange between Cdc42 and Par-3. Interestingly, in the structure of the Par complex with the substrate Lethal giant larvae (Lgl), Lgl binds the Par-6 PDZ in a manner that precludes its interaction with the aPKC kinase domain, suggesting that it may disrupt this interaction (42). Future efforts will be directed at understanding how the interactions might alter other Par complex functions, such as aPKC catalytic activity and membrane recruitment.

## Experimental Procedures

## Data availability

All data are contained within the manuscript.

### Cloning

GST-, MBP- and his-tagged constructs were cloned as previously described (13, 20) using Gibson cloning (New England BioLabs), Q5 mutagenesis (New England BioLabs) or traditional methods. In addition to an N-terminal MBP tag, the aPKC PB1-C1 (residues 1-225) construct also contained a C-terminal his-tag. Par complex components (aPKC and his-Par-6) were cloned into pCMV as previously described (20, 39). Please see the Key Resources table for additional information on specific constructs.

### Expression

Par complex, full length aPKC and full length aPKC with Par-6 PB1 were expressed in HEK 293F cells (Thermofisher), as previously described (20, 39). Briefly, cells were grown in FreeStyle 293 expression media (Thermofisher) in shaker flasks at 37°C with 8% CO2. Cells were transfected with 293fectin (Thermofisher) or ExpiFectamine (Thermofisher) according to the manufacturer’s protocol. After 48 hours, cells were collected by centrifugation (500g for 5 min). Cell pellets were resuspended in nickel lysis buffer [50mM NaH2PO4, 300 mM NaCl, 10 mM Imidazole, pH 8.0] and then frozen in liquid N2 and stored at -80°C.

All other proteins were expressed in E. coli (strain BL21 DE3). Constructs were transformed into BL21 cells, grown overnight at 37°C on LB + ampicillin (Amp; 100 mg/mL). Resulting colonies were selected and used to inoculate 100mL of LB + Amp starter cultures. Cultures were grown at 37°C to an OD600 of 0.6-1.0 and then diluted into 2L LB + Amp cultures. At an OD600 of 0.8-1.0 expression was induced with 0.5 mM IPTG for 2-3 hours. Cultures were centrifuged at 4400g for 15 minutes to pellet cells. Media was removed and pellets were resuspended in nickel lysis buffer [50mM NaH2PO4, 300 mM NaCl, 10 mM Imidazole, pH 8.0], GST lysis buffer [1XPBS, 1 mM DTT, pH 7.5] or Maltose lysis buffer [20 mM Tris, 200 mM NaCl, 1 mM EDTA, 1 mM DTT, pH 7.5], as appropriate. Resuspended pellets were frozen in liquid N2 and stored at -80°C.

### Purification

Resuspended E.coli pellets were thawed and cells were lysed by probe sonication using a Sonicator Dismembrator (Model 500, Fisher Scientific; 70% amplitude, 0.3/0.7s on/off pulse, 3×1 min). 293F cell pellets were lysed similarly using a microtip probe (70% amplitude, 0.3/0.7s on/off pulse, 4×1 min). Lysates were centrifuged at 27,000g for 20 min to pellet cellular debris. GST-tagged protein lysates were aliquoted, frozen in liquid N2 and stored at -80°C.

His-tagged protein lysates, except for aPKC KD-PBM and KDΔPBM, were incubated with HisPur Cobalt (Thermofisher) resin for 30 min at 4°C and then washed 3x with nickel lysis buffer. For 293F lysates, 100µM ATP and 5mM MgCl2 were added to the first and second washes. Proteins were eluted in 0.5-1.5mL fractions with nickel elution buffer [50 mM NaH2PO4, 300 mM NaCl, 300 mM Imidazole, pH 8.0]. For all proteins expressed in E.coli, fractions containing protein were pooled, buffered exchanged into final buffer [20mM HEPES pH 7.5, 100 mM NaCl and 1 mM DTT] using a PD10 desalting column (Cytiva), concentrated using a Vivaspin20 protein concentrator spin column (Cytiva), aliquoted, frozen in liquid N2 and stored at -80°C. For 293F-expressed constructs, proteins were further purified using anion exchange chromatography on an AKTA FPLC protein purification system (Amersham Biosciences). Following his-purification fractions were pooled and buffered exchanged into 20mM HEPES pH 7.5, 100 mM NaCl, 1 mM DTT, 100 µM ATP and 5 mM MgCl2 using a PD10 desalting column (Cytiva). Buffer-shifted protein was injected onto a Source Q (Cytiva) column and eluted over a salt gradient of 100-550mM NaCl. Fractions containing the desired protein(s) were pooled, buffered exchanged into 20 mM HEPES pH 7.5, 100 mM NaCl, 1 mM DTT, 100 µM ATP, and 5 mM MgCl2 using a PD10 desalting column (Cytiva), concentrated using a Vivaspin20 protein concentrator spin column (Cytiva), aliquoted, frozen in liquid N2 and stored at -80°C.

Due to solubility issues, aPKC KD-PBM and KDΔPBM were expressed in E. coli and his-purified partially under denaturing conditions. Following sonication and centrifugation (described above), the soluble fraction was discarded and the insoluble pellet was resuspended in 50mM NaH2PO4, 300 mM NaCl, 10 mM Imidazole, 8M Urea pH 8.0. Centrifugation was repeated (27,000g for 20 min) and the resulting soluble phase was incubated with HisPur Ni-NTA resin (ThermoFisher) for 30 min at 4°C. Resin was washed and eluted as described above. Purified protein was aliquoted, frozen in liquid N2 and stored at -80°C.

### Qualitative binding assays

For qualitative binding (“GST pulldown”) assays, GST lysates were incubated with glutathione agarose resin (GoldBio) for at least 30 min at 4°C and then washed 3x 5 min washes at room temp with binding buffer [10 mM HEPES pH 7.5, 100 mM NaCl, 1 mM DTT 200 µM ATP, 5 mM MgCl2 and 0.1% Tween-20] with rotational mixing. Soluble proteins were added to GST-bound proteins, as indicated, and incubated at room temperature with rotational mixing for 60 min. Resin was then washed 3x with binding buffer and proteins were eluted with 4X LDS sample buffer (ThermoFisher). Samples were run on a Bis-Tris gel and stained with Coomassie Brilliant Blue R-250 (GolBio). Band intensities of replicates were quantified using ImageJ (v1.53a). The normalized band intensity was determined by averaging the intensity of either full length aPKC or KD-PBM signal within the experiment, as appropriate, and then dividing the band intensity of each individual value by that mean value. The data was visualized and analyzed using the DABEST (43) software packages. Confidence intervals were estimated using the bootstrap method as implemented in DABEST.

## Acknowledgments

This work was supported by NIH grants R35GM127092 and T32HD007348.

## Author Contributions

E.V., R.R.P. and K.E.P. designed the experiments, analyzed the data, prepared figures and wrote the manuscript. E.V. and R.R.P. performed the experiments.

